# Salmonella SPI2 evades detection by NAIP/NLRC4 despite utilization of a detectable needle

**DOI:** 10.64898/2026.01.05.696446

**Authors:** Jinyi (Sammy) Guo, Farhan K. Lakhani, Cody G. Vientos, Lupeng Li, Edward A. Miao

## Abstract

*Salmonella enterica* serovar Typhimurium uses two type 3 secretion (T3S) systems to reprogram host cells. Whereas activity of the SPI1 T3S can be detected by the NAIP/NLRC4 inflammasome, the SPI2 T3S is not detected. In this study, we demonstrate that the SPI2 needle protein, SsaG, can be detected by the NAIP-NLRC4 inflammasome when artificially delivered to the cytosol in mouse macrophages. Surprisingly, this detection occurs primarily through NAIP2, which normally detect rod proteins. However, we found that during infection with live *S.* Typhimurium, this detection fails to trigger the downstream pyroptotic pathway. We investigate the hypothesis that the SPI2 effector protein SpvC inhibits NLRC4 activation. In mouse macrophages, *spvC* mutants were detected under SPI2-inducing conditions in an NLRC4-dependent manner. However, this masking effect of SpvC is not seen *in vivo*, where SpvC contributes to *S.* Typhimurium virulence in mice independent from NLRC4. We also hypothesized that *S.* Typhimurium may minimize SsaG detection by limiting protein expression. In support of this, overexpressing SsaG causes limited NLRC4-dependent clearance in vivo. Therefore, the SPI2 may evade NAIP/NLRC4 inflammasome detection by precisely controlling the quantity of detectable SsaG needle protein expressed.

## Introduction

The host innate immune system serves as the first line of defense against infection. This defense relies on a network of cytosolic pattern recognition receptors (PRRs) that can be activated by specific bacterial ligands, initiating a range of host responses. A critical component of this PRR network is the nucleotide-binding domain, leucine-rich repeat-containing (NLR) family, about a third of which form multi-protein inflammasome complexes^1^. Among these, the NAIP-NLRC4 inflammasome is a key sensor for Gram-negative bacteria, specialized to detect the activity of the bacterial type III secretion system (T3SS), a syringe-like apparatus used to inject virulence proteins from the bacterial cytosol directly into the cytosol of host cells. These effectors reprogram host cell physiology in ways that benefit the pathogen.

T3SS effectors are highly variable between different pathogen species, and therefore detection of any effector requires a specific sensor. Mammals do use specific sensors to detect some effectors; for example, the pyrin inflammasome detects the activity of the *Yersinia* YopE and YopT effectors^2^. This strategy is used by plants, which encode hundreds of NLR-like sensors, many of which detect the activity of individual effectors^3,4^. In vertebrates, a more general detection strategy is also used that capitalizes upon errors in the T3SS injection process. Although the T3SS evolved to inject effectors, it also inadvertently injects small amounts of flagellin, T3SS rod, and T3SS needle proteins. These highly conserved proteins are detected by the NAIP/NLRC4 inflammasome^5,6,7,8,9,10^.

The three ligands are detected by direct binding to specific NAIP subfamily of sensors. NAIPs were originally named as “NLR family, apoptosis inhibitory proteins”, although we now know these are not related to apoptosis or inhibition. Although NAIPs exhibit significant species-specific differences, their overall function between species is conserved^9^. In mice, an expanded family of NAIPs provides specific detection of the three ligands: NAIP1 recognizes the T3SS needle protein^9,10^, NAIP2 detects the rod protein^7,8^, and NAIP5/6 both recognize flagellin^510^. In contrast, humans encode only one functional NAIP, which senses the T3SS rod^11^, needle^9^, and flagellin^12^ via one single isoform^11^. Upon binding a ligand, NAIPs recruit NLRC4 (NLR family CARD domain containing 4), triggering its oligomerization into a ring-like "inflammasome" structure^8,13,14^. This inflammasome complex activates caspase-1, which in turn cleaves pro-inflammatory cytokines interleukin 1 beta (IL-1β) and interleukin-18 (IL-18) and the pore-forming protein Gasdermin D (GSDMD) to their active forms^15,16,17,18,19^. Cleaved GSDMD forms large pores that lead to a lytic, inflammatory cell death known as pyroptosis.^20,21^ Pyroptosis is crucial for controlling bacterial infections and is especially effective in eliminating the intracellular niche of environmental pathogens that fail to evade inflammasome detection^22,23,24^.

*Salmonella enterica* serovar Typhimurium (*S.* Typhimurium) is a formidable pathogen that masterfully exploits host cell processes using two distinct T3SSs. The *Salmonella* Pathogenicity Island 1 (SPI1) T3SS is required for the initial invasion of non-phagocytic cells, allowing the bacterium to gain access to a vacuolar compartment of intestinal epithelial cells^25,26^. The SPI2 T3SS is expressed after internalization where it translocates effector proteins that facilitate *Salmonella*’s survival and replication within the *Salmonella*-containing vacuole (SCV). SPI2 is critical for intracellular vacuolar replication in both intestinal epithelial cells^27^ and macrophages^26,27,28^ the two primary cell-type niches that support *Salmonella* replication.

Within this two-phase lifestyle lies a paradox in *Salmonella* pathogenesis^29^, it is well-established that the SPI1 T3SS rod (PrgJ) and needle (PrgI) proteins are potent ligands for the NAIP-NLRC4 inflammasome. For mouse macrophages, priming with poly(I:C) for two days induces NAIP1 expression and conferred sensitivity to PrgI^10^. Furthermore, flagellin is expressed in the gut lumen in the same environments where *Salmonella* expresses SPI1. However, the SPI2 T3SS, which is essential for replication in macrophages, appears to evade this detection.

This necessity for stealth is best exemplified by the SPI2 rod protein, SsaI. Although SsaI shares 33.7% sequence similarity with its SPI1 homolog PrgJ, it is not detected by NAIP2 in mouse macrophages^7^. This evasion is driven by specific evolutionary adaptations: SsaI possesses a distinct C-terminal sequence compared to PrgJ.

*Salmonella* employs a multi-layered strategy to hide its flagellin. While the SPI1 flagellin (FliC) is a potent NAIP5/6 ligand, *Salmonella* avoids flagellin detection during the intracellular phase primarily through transcriptional repression^30^. This evasion is critical, as *S.* Typhimurium strains that are experimentally engineered to deprive them of inflammasome activation leads to rapid pyroptotic death of the host macrophage, prematurely ending the bacterium’s replicative niche. When *Salmonella* is engineered to constitutively express FliC under SPI2 conditions ("FliC-ON"), the bacteria are immediately detected by NLRC4 and cleared^7^. The alternate phase 2 flagellin (FljB) can also activate the NLRC4 inflammasome^7^. The same strong attenuation is observed when the SPI1 rod protein is expressed under a SPI2 promoter (PrgJ-ON)^7^. These data strongly support the paradigm that SPI2 phase proteins have evolved to be invisible, at least for the SsaI rod and flagellin.

SPI2 also encodes a needle protein, SsaG. Because SsaG functions similarly to the SPI1 needle PrgI, sharing 32.1% sequence similarity^7^, and operates in the stealthy SPI2 phase, we previously assumed it would similarly evade detection. However, previous work found that SsaG can be detected by human macrophages (THP-1 cells)^31^. Here, we test whether SsaG can be detected in mouse macrophages.

## Results

### Cytosolic SPI2 needle protein SsaG activates the murine NLRC4 inflammasome

Previous studies have established that the *Salmonella* SPI1 T3SS rod protein (PrgJ) is a potent ligand for the murine NAIP-NLRC4 inflammasome, whereas the homologous SPI2 rod protein (SsaI) evades detection. Given the structural conservation between the two secretion systems, we investigated whether the SPI2 needle protein, SsaG, possesses ligand properties capable of activating NLRC4 if exposed to the host cytosol.

To test this, we utilized a retroviral transduction system to ectopically express T3SS components in immortalized bone marrow-derived macrophages (iBMMs). We cloned the genes encoding the SPI1 rod (*prgJ*) and needle (*prgI*), as well as the SPI2 rod (*ssaI*) and needle (*ssaG*), into a GFP-expressing retroviral vector (pMXs-IG). In this reporter assay, if the transduced proteins do not trigger pyroptosis, then GFP-positive cells are observed, which is quantified via flow cytometry. In contrast, ligands that activate the inflammasome cause pyroptosis, and therefore all transduced cells are deleted from the assay which causes the loss of GFP-positive transduced cells.

Transduction of the known SPI1 agonist PrgJ resulted in a significant reduction of GFP-positive cells in wild-type (WT) iBMMs compared to *Nlrc4*^−/−^ controls, validating the assay (**Fig. 1A, B**). Strikingly, transduction of the SPI2 needle protein SsaG triggered a similar, robust loss of GFP-positive cells in a strictly NLRC4-dependent manner (**Fig. 1A, B**). Conversely, neither the SPI1 needle protein (PrgI), nor the SPI2 rod protein (SsaI) induced significant pyroptosis in this system (**Fig. 1A, B**). The lack of PrgI detection is consistent with the requirement for intensive host priming for NAIP1 expression that was not used in the experiment here. The lack of SsaI responsiveness confirms its evasion of NAIP2 detection. Note that these cells were not primed with TLR ligands.

**Figure 1:**
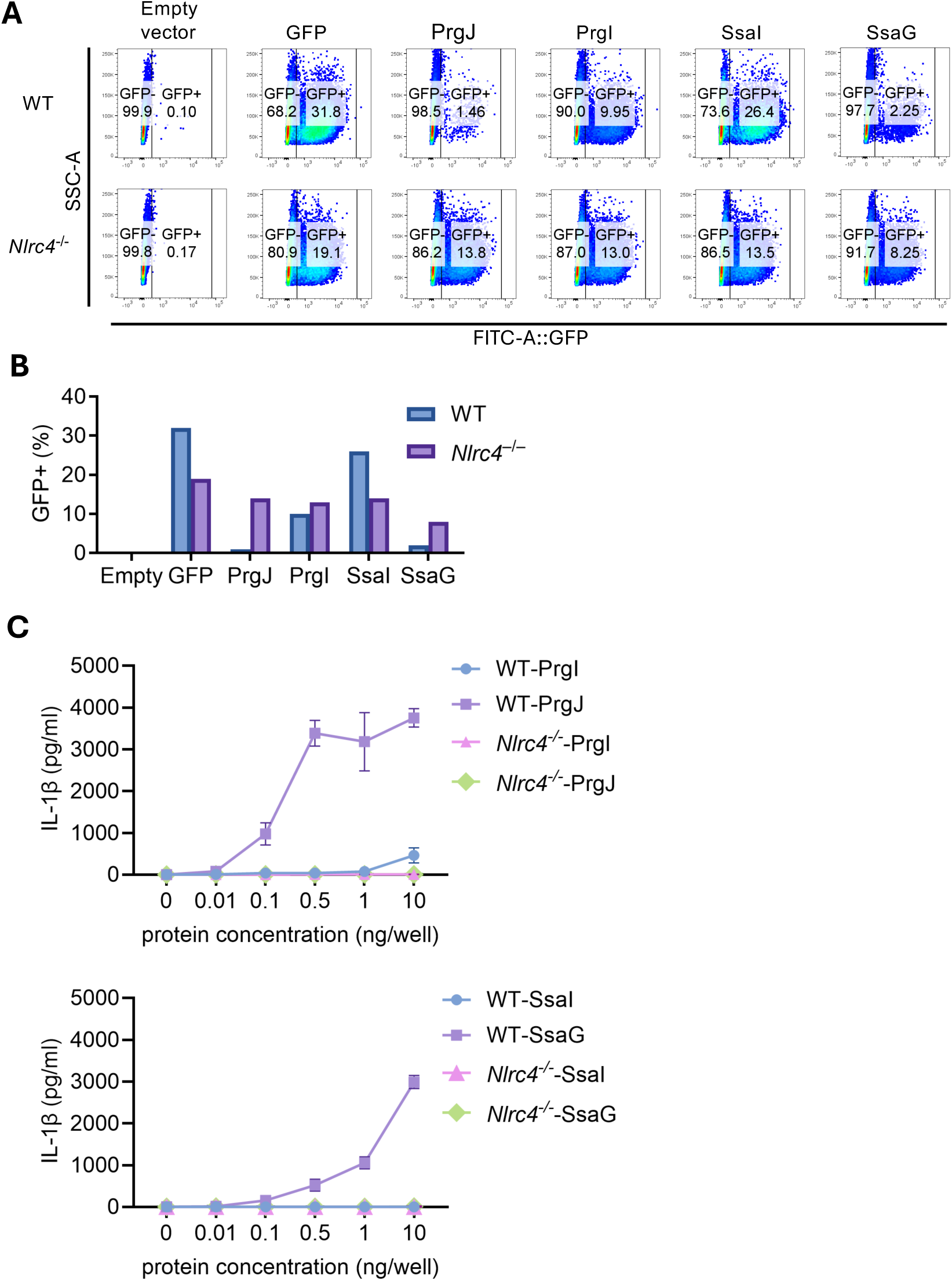
Cytosolic SPI-2 needle protein SsaG activates the murine NLRC4 inflammasome. (A) iBMM were infected with transgenic retroviruses expressing GFP alone, or prgJ, prgI, ssaI, ssaG, respectively, followed by an IRES-GFP element. GFP positive cells were identified by flow cytometry 2 days after infection. (B) Quantitation of (A) (C) Indicated quantity of purified protein was delivered to Pam3CSK4 primed BMM by protein transfection, and IL-1β secretion was determined by ELISA.

To corroborate these findings, we purified each SPI1 and SPI2 rod and needle proteins as His-tagged fusion proteins. We transfected LPS-primed macrophages with the purified proteins and then measured IL-1β secretion as an inflammasome-dependent readout. Consistent with the retroviral assay, the cytosolic delivery of SsaG induced robust IL-1β secretion, though higher amounts of SsaG were required to achieve levels comparable to those triggered by PrgJ positive control (**Fig. 1C**). Meanwhile, PrgI detection was minimal, consistent with the inadequate NAIP1 expression (again these cells have not been exposed to prolonged priming)^10^. Further, SsaI was again not detected. This response was abrogated in *Nlrc4*^−/−^ macrophages, confirming that cytosolic SsaG can activate the murine NLRC4 inflammasome. These results were surprising given that PrgI and SsaG function as homologous needle subunits within SPI1 and SPI2 T3SS apparatuses, respectively. PrgI is detected by NAIP1 only after prolonged poly(I:C) priming in BMMs^910^; in contrast, SsaG detection was efficient, suggesting that it is detected by additional NAIPs besides NAIP1. Therefore, we next investigate which NAIP is responsible to detect SsaG.

### SsaG detection is partially mediated by NAIP2

Murine NAIPs exhibit ligand specificity, with NAIP1 and NAIP2 directly bind the T3SS needle and rod, respectively^4,8,9,32^. To more specifically determine which NAIP contributes to SsaG recognition, we transfected the SsaG protein into a panel of macrophages deficient in individual NAIPs (*Naip1*, *Naip2*, *Naip5*) or functional deficient of the entire cluster (*Naip1-6*).

SsaG-induced inflammasome activation remained intact in *Naip1*^−/−^ and *Naip5*^−/−^macrophages, exhibiting levels comparable to WT macrophages (**Fig. 2A**). In contrast, activation was significantly reduced in *Naip2*^−/−^ macrophages, although a residual detection was observed (**Fig. 2A, B**). As expected, inflammasome activation was completely abolished in *Naip1-6*^−/−^ cells. These data indicate that NAIP2—classically defined as the rod sensor—can detect the SPI2 needle SsaG. However, it is not the sole receptor, suggesting functional redundancy with another NAIP paralog.

**Figure 2:**
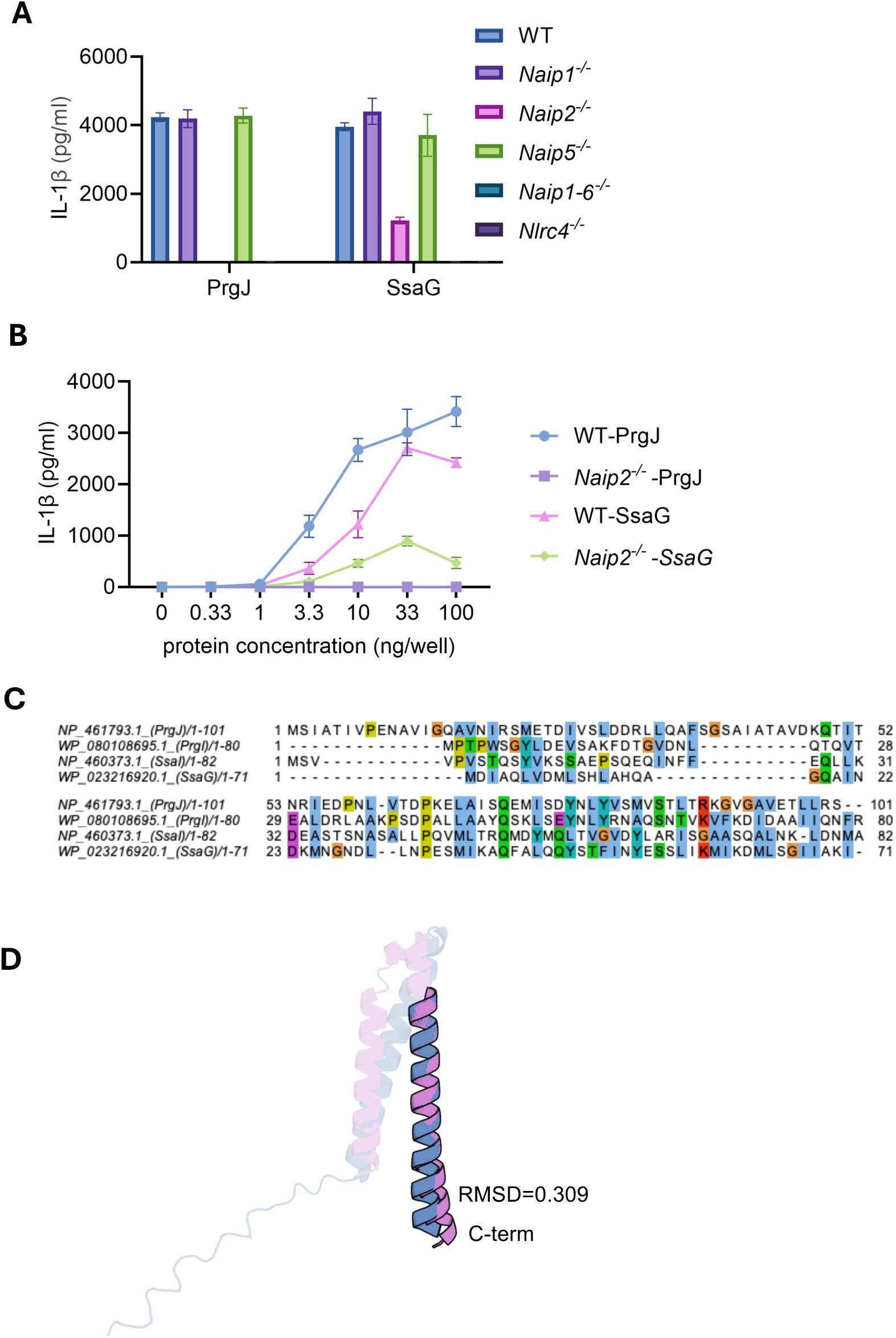
SsaG detection is partially mediated by NAIP2. (A) A 10ng quantity of indicated purified protein was delivered to Pam3CSK4 primed BMM by protein transfection, and IL-1β secretion was determined by ELISA. (B) Indicated quantity of purified protein was delivered to Pam3CSK4 primed WT or *Naip2*^-/-^ BMM by protein transfection, and IL-1β secretion was determined. (C) Protein sequence alignment. MEGA12 was used for alignment, and clusters were determined with Jalview. (D) Structure of PrgJ and SsaG with C-terminus aligned. Blue indicates PrgJ and pink indicates SsaG.

The observation that SsaG detection is partially mediated by NAIP2 is unexpected, as NAIP2 is the canonical sensor for the T3SS rod protein (PrgJ), whereas the T3SS needle protein (PrgI) is detected by NAIP1. Structurally, SsaG is the homolog of the SPI1 needle PrgJ (**Fig. 2C**). They share the specific helical architecture required for needle polymerization and approximately 32.1% sequence similarity. However, SsaG also shares 22.5% similarity with the SPI1 rod, PrgJ^7^. While sequence alignment provides limited insight, we next compared the predicted tertiary structures of SsaG and PrgJ (**Fig. 2D**). The global RMSD is 6.502 Å, indicating the overall structures are distinct. However, alignment of the C-termini reveals a local RMSD of 0.309Å, indicating these specific domain structures are highly similar. Since mouse NAIP2 is known to bind the C-terminus of PrgJ^33^, these data suggest that NAIP2 recognition is not strictly limited to rod proteins, but can cross-react with the SPI2 needle due to shared structural features.

### SpvC inhibits NLRC4 inflammasome detection in vitro but contributes to virulence via an NLRC4-independent mechanism in vivo

Our finding that cytosolic SsaG is a potent ligand for the NAIP-NLRC4 inflammasome stands in sharp contrast to the established observation that live, SPI2-expressing *Salmonella* evade this detection. This discrepancy suggests that *Salmonella* might employ an active mechanism to suppress SsaG sensing during macrophage infection. A primary candidate for this inhibitory role is SpvC, a known SPI2 T3SS effector protein and phosphothreonine lyase that irreversibly inactivates host MAPK signaling pathways, including pERK, pJNK, and p38. Previous studies have suggested that SpvC can suppress pyroptosis^34^, inhibit autophagy^35^, or undermines neutrophil defense^36^. A significant clue to the potential function of SpvC comes from *Shigella*, where the effector OspF, a homolog of SpvC, was recently shown to inhibit NLRC4 inflammasome^37^. Given the shared homology and enzymatic function of SpvC and OspF, we hypothesized that SpvC performs an analogous role in *Salmonella*, actively masking the detection of the SPI2 T3SS from the NLRC4 inflammasome.

To test this, we generated a Δ*spvC* mutant *S.* Typhimurium strain by lambda-Red mediated homologous recombination. We then infecsted primary mouse BMMs with wild-type (WT) *S.* Typhimurium or the isogenic Δ*spvC* mutant. To ensure the bacteria were in the appropriate physiological state, strains were sub-cultured in SPI2-inducing media to mimic the vacuolar environment prior to infection. In agreement with previous literature, WT *Salmonella* induced only minimal inflammasome activation; however, infection with the Δ*spvC* mutant resulted in significantly higher and more rapid activation (**Fig. 3A**). FliC^ON^ *S.* Typhimurium were used as a positive control strain that activates NLRC4 when grown under SPI2-inducing conditions. This enhanced activation was completely abrogated in *Nlrc4*^−/−^ macrophages, demonstrating that SpvC is essential for evading NLRC4-mediated detection during macrophage infection.

**Figure 3:**
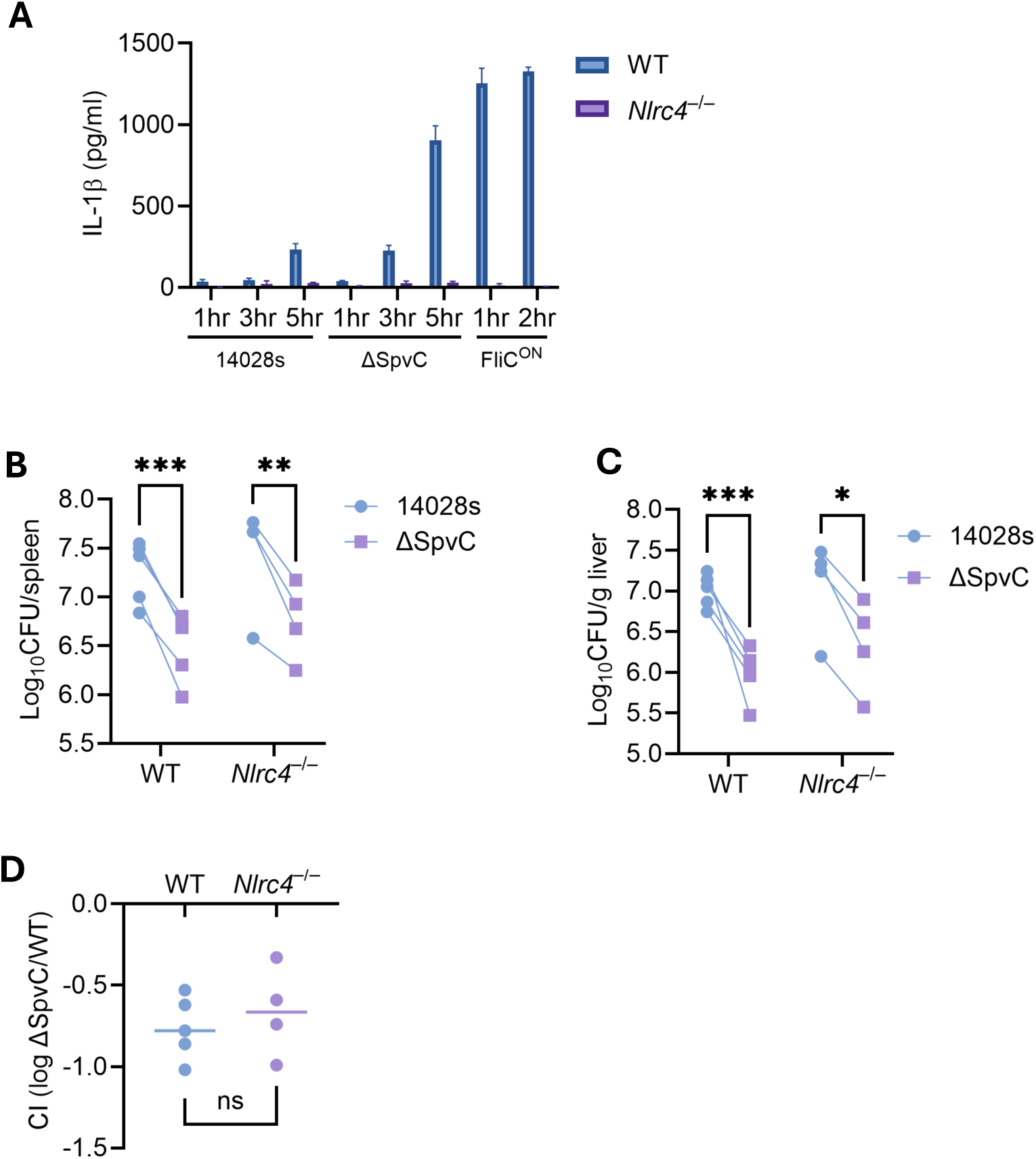
SpvC inhibits NLRC4 inflammasome detection in vitro but contributes to virulence via an NLRC4-independent mechanism in vivo. (A) BMMs were infected with indicated SPI-2 induced *S.*Typhimurium strains. Supernatant were collected and IL-1β secretion was determined by ELISA. (B-D) Mice were infected with a 1:1 ratio of Δ*spvC* and a vector control *S.*Typhimurium strain. Bacterial burdens in the spleen and liver were determined at 4 days post infection. Ratio of the mutant Δ*spvC* and vector control is graphed. Two-way repeated measure ANOVA performed to determine difference between the competitive index of these two strains in different mice.

Therefore, we hypothesized that the primary *in vivo* role of SpvC is to suppress the NLRC4 inflammasome. We first confirmed SpvC plays the role as a virulence factor using single-strain infections in C57BL/6J WT mice, where the Δ*spvC* mutant showed a significantly lower bacterial burden in the spleen and liver compared to WT bacteria (**Fig. 3B, C**). To determine if this virulence defect was mediated by unmasked NLRC4 detection, we performed a competitive index (CI) assay by co-infecting mice with WT and Δ*spvC* strains. Each strain encoded a different antibiotic resistance, allowing us to quantify the number of bacteria from each strain within the same host. As expected, the Δ*spvC* mutant was outcompeted in WT mice, resulting in a negative competitive index (**Fig. 3D**). However, contradicting our hypothesis, the Δ*spvC* mutant was outcompeted to the same degree in *Nlrc4*^−/−^ mice (**Fig. 3D**). This result demonstrates that while SpvC is a critical virulence factor in vivo, its primary contribution to *in vivo* fitness is independent of NLRC4.

### Overexpression of SsaG drives inflammasome-mediated clearance independent of SpvC *in vivo*

Our previous data established that: 1) cytosolic SsaG is a ligand for NLRC4, yet 2) *in vivo* evasion of this detection is SpvC-independent. Another piece of evidence arguing against SpvC as a penetrant NLRC4 inhibitor is that FliC^ON^ and PrgJ^ON^ bacteria are readily detected, despite their ability to express SpvC^7^. Indeed, the mechanism by which the SpvC homolog OspF inhibits NLRC4 activity during *Shigella* infection is by inhibiting a rapid priming effect that is required in human macrophages, which seems not required in mouse macrophages^37^. We next hypothesized that the natural levels of SsaG translocated during infection are simply too low to trigger the inflammasome *in vivo*. We reasoned that if we artificially increased the intracellular concentration of SsaG in an engineered strain, it might then cross the threshold for detection.

To test this, we engineered a strain that constitutively overexpresses the SPI2 needle protein (SsaG^ON^) in WT *Salmonella*. We infected WT and *Casp1/11*^−/−^ mice and compared the bacterial burdens to WT *Salmonella*. In contrast to natural infection levels, overexpression of SsaG resulted in significantly lower bacterial burdens compared to the parent strain in WT mice (**Fig. 4A, B**). Crucially, this clearance phenotype was completely abrogated in *Casp1/11*^−/−^ mice, confirming that the high levels of SsaG were triggering a caspase-1/11-dependent inflammasome response (**Fig. 4A, B**). This demonstrates that the *in vivo* immune system is capable of detecting SsaG in the context of SPI2-expressing *S.* Typhimurium infection in vivo, provided the ligand is abundant enough. However, we do note that the magnitude of clearance of SsaG^ON^ is much lower than that observed for FliC^ON^ and PrgJ^ON^ ^7,38^.

**Figure 4:**
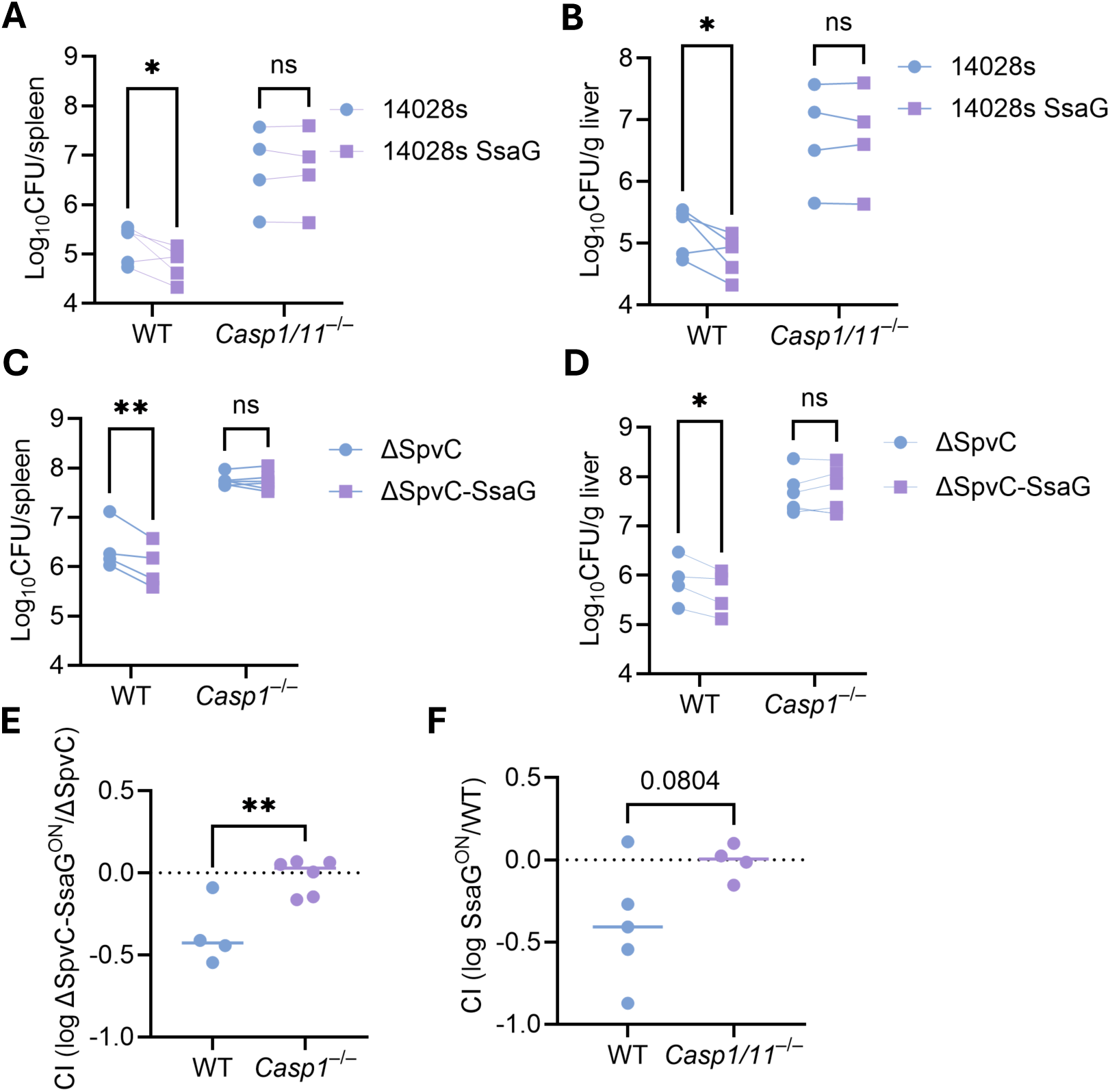
Overexpression of SsaG drives inflammasome-mediated clearance independent of SpvC. (A-F) Mice were infected with a 1:1 ratio of indicating *S.*Typhimurium strains. Bacterial burdens in the spleen and liver were determined at 4 days post infection. Ratio of the mutant strain and its respective vector control is graphed. Two-way repeated measure ANOVA performed to determine difference between the competitive index of these two strains in different mice.

We next asked if the presence of the SpvC can inhibit inflammasome detection against this high-abundance SsaG signal provided by SsaG^ON^. We expressed the SsaG^ON^ construct in the Δ*spvC* mutant strains. If SpvC functions as a stoichiometric inhibitor, we expect it to dampen the detection of SsaG. However, the Δ*spvC*-SsaG^ON^ strain was cleared significantly better than the Δ*spvC* parent strain in WT mice, a defect that was rescued in *Casp1*^−/−^mice (**Fig. 4C, D**).

We compared the magnitude of the clearance in both backgrounds. We found that the Competitive Index (CI) between the Δ*spvC* strains (Δ*spvC* vs. Δ*spvC*-SsaG^ON^) was statistically indistinguishable from the CI between the WT strains (WT vs. WT-SsaG^ON^) (**Fig. 4E, F**). This indicates that the detection of SsaG overexpression is identical whether SpvC is present or not. Together, these data suggest that while SsaG is a bona fide ligand for the NLRC4 inflammasome, SpvC is unable to mask it when the ligand exceeds its normal protein expression threshold, further supporting the conclusion that the primary *in vivo* function of SpvC may lie outside the NLRC4 axis.

## Discussion

The SPI2 T3S system of *Salmonella* is traditionally viewed as immunologically silent, in sharp contrast to the SPI1 system which triggers robust detection by the NLRC4 inflammasome. It was previously shown that SPI2 needle protein SsaG can be detected by human THP-1 cells^31^. Our work identified SsaG as an effective ligand for the murine NLRC4 inflammasome, which complicate the understanding of this stealthy SPI2 system. Even more striking is that the sensor responsible for this detection is NAIP2, which is normally assigned to rod proteins, rather than NAIP1, the canonical sensor of the needle.

This sensor mismatch argues that murine NAIP proteins rely on recognizing structural features rather than strictly monitoring primary sequence. Although SsaG shares only about 22.5% similarity with the SPI1 rod protein PrgJ^7^, both proteins must assemble into a conserved helical scaffold to stack within the secretion apparatus. We propose that NAIP2 detection of the SPI2 needle arises from this shared architectural constraint because of the structural similarity of their C-terminal domains. This suggests that *Salmonella* can alter sequence to reduce visibility to the immune system, but only up to a point. The fundamental geometric requirements of T3SS assembly force SsaG to retain a C-terminal structural motif that resembles PrgJ, allowing NAIP2 to bind. Thus, structural necessity traps the pathogen in an evolutionary trade-off: the needle cannot escape recognition without compromising the integrity of the secretion apparatus that is essential for virulence.

The most surprising outcome of our work is the difference between *in vitro* and *in vivo* responses. *In vitro*, the SsaG-triggered pyroptotic response requires inhibition by SpvC, implying that SsaG can be strongly detected and that the detection depends on SpvC. *In vivo*, however, the virulence defect of the Δ*spvC* mutant is independent of NLRC4, even though NLRC4 can detect SsaG. One explanation is ligand abundance. For instance, previous evidence showed that PrgJ^ON^ can be readily detected when it is overexpressed^7^, which naturally is not expressed in SPI2 condition.

Our *in vitro* model, which relies on SPI2 pre-induction, may introduce an experimental artifact by loading the bacteria with supraphysiological levels of SsaG prior to macrophage contact. This likely creates a bolus release of ligand that exceeds the host’s detection threshold, a scenario unlikely to occur during natural infection. Indeed, our SsaG overexpression data suggest that *Salmonella* normally evades detection by tightly repressing SsaG transcription to remain below this activation threshold. However, ligand abundance appears to be only part of the equation. We observed that even when overexpressing SsaG, the resulting clearance is markedly less efficient than that triggered by flagellin (FliC-ON) or the SPI1 rod (PrgJ-ON)^7^. This discrepancy suggests a secondary layer of evasion linked to the structural properties of the protein itself. Recent study also showed that fusing N-terminus of YopE with C-terminus of either SsaI or SsaG to facilitate T3SS ligand translocation can also allow these ligands to be detectable upon priming^39^. One explanation is that these fusion protein to YopE changes the folding in these proteins somewhat that makes it detectable, suggesting that the evasion is more complex than simply a few amino acid changes. It is therefore plausible that native SsaG and SsaI adopt a monomeric fold that masks their immunogenic motifs, a stealth conformation that limits detection even when cytosolic delivery is successful. Thus, we propose that *Salmonella* employs a composite evasion strategy: primarily relying on transcriptional repression to limit ligand quantity, while simultaneously benefiting from an intrinsic structural defect that raises the threshold required for NAIP2 activation.

If SpvC is not required to shield the bacterium from inflammasome-mediated killing *in vivo*, what drives the attenuation of the Δ*spvC* mutant? We propose that SpvC might interfere with a central regulator of host inflammatory stress. The MAPK pathways, particularly p38 and JNK, act as critical integration hubs that translate bacterial detection into a high-alert inflammatory state, driving the transcription of potent cytokines (e.g., TNF and IL-6)^40^ and regulating downstream cell fate decisions. In the absence of SpvC, the Δ*spvC* mutant likely triggers unrestricted MAPK signaling, resulting in a hyper-inflammatory environment that accelerates bacterial clearance. By deploying SpvC to irreversibly dephosphorylate these kinases, *Salmonella* might sever these signaling pathways. This blockade can prevent the macrophage from mounting a coordinated cytokine response, thereby preserving the intracellular niche against the broader immune recruitment that would otherwise eliminate the infection.

Taken together, our findings redefine the SPI-2 needle SsaG as a conditionally detectable ligand recognized by the NAIP2-NLRC4 inflammasome. We propose that Salmonella mitigates this structural liability through tight transcriptional repression to maintain SsaG levels below the activation threshold while deploying SpvC to neutralize MAPK-driven inflammatory priming. The complete absence of NLRC4 activation during wild-type infection suggests a critical physical mechanism prevents ligand access to the sensor. Future studies must determine whether SsaG evasion is driven by a failure to translocate from the vacuole or if cytosolic SsaG is subject to rapid neutralization. Elucidating whether the mechanism relies on physical sequestration or active silencing will define the precise strategy intracellular pathogens use to operate essential secretion machinery without alerting the host.

## Materials and Methods

### Mice and Cell Culture

Wild-type C57BL/6 and *Nlrc4*-deficient (*Nlrc4*^−/−^) mice^38^ were housed in a specific pathogen–free facility. All animal experiments were performed under the supervision and approval of the Institutional Animal Care and Use Committee (IACUC). Bone marrow-derived macrophages were differentiated from femoral bone marrow precursors by culturing in complete DMEM media (DMEM+10% FBS) supplemented with L-cell conditioned supernatants (15%)^38^. For retroviral experiments, Phoenix-Ampho retroviral packaging cells (American Type Culture Collection) and immortalized bone marrow-derived macrophages were maintained in complete DMEM media.

### Bacterial Strains and Growth Conditions

*Salmonella enterica* serovar Typhimurium wild-type, ssaG-ON, and spvC mutant strains were used in this study. The spvC deletion mutant was constructed using Lambda Red recombineering^41^. For *in vitro* infection assays, bacterial cultures were grown under conditions that induce *Salmonella* Pathogenicity Island-2 (SPI2) expression. Briefly, overnight cultures were back-diluted to an optical density at 600 nm (OD₆₀₀) of 0.026 in 3 mL of SPI2 inducing media and grown for 16–20 hours in a shaking incubator at 37°C.

### Retroviral Transduction

Retroviral vectors pMXsIP encoding mIpaf-PC (Protein C epitope tag) or PC-GFP were generated by transfecting Phoenix-Ampho cells using Lipofectamine 2000 (Invitrogen) according to the manufacturer’s instructions. The resulting retroviral supernatants were harvested and used to transduce iBMDMs. Transduced populations were selected with puromycin, and expression efficiency was confirmed by flow cytometry.

### Protein Transfection

BMDMs were seeded in 96-well plates at a density of 1 × 10⁵ cells per well. Cells were primed with 500 ng/mL Pam3CSK4 for 3 hours prior to transfection. Purified proteins were transfected into macrophages using the Profect P2 transfection reagent. Unless otherwise indicated, proteins were transfected at a concentration of 10 ng/well for 1 hour.

### Macrophage Infection Assays

For bacterial infections, BMDMs were seeded at a density of 5 × 10⁴ cells per well and primed with lipopolysaccharide (LPS; 10 ng/mL) for 16 hours prior to infection. Cells were infected with SPI2 induced^42^ *S*. Typhimurium strains at a multiplicity of infection (MOI) of 10. One hour post-infection, extracellular bacteria were eliminated by treating cells with media containing gentamicin (50 µg/mL).

### ELISA

Cell culture supernatants were harvested at the indicated time points post-infection and clarified by centrifugation to remove cell debris and bacteria. The concentration of secreted IL-1β was quantified using a mouse IL-1β DuoSet ELISA kit (R&D Systems, Catalog #DY401) according to the manufacturer’s instructions. Optical density (OD) was measured at 570 nm using a microplate reader, and concentrations were calculated based on a standard curve.

### Competitive Index and Systemic Infections

The competitive index (CI) experiments were performed as previously described^38^. Briefly, the inoculum was prepared by mixing the vector control strain and the experimental strain at a 1:1 ratio. Mice were infected intraperitoneally with a total dose of 1×10^5^ CFU of *S*. Typhimurium. At 4 days post-infection, spleens and livers were harvested and homogenized in 2 mL tubes containing 1 mL of sterile PBS and a single 5 mm stainless steel bead. Organ homogenates were serially diluted in sterile PBS and plated on Luria-Bertani agar plates supplemented with appropriate selective antibiotics to enumerate colony-forming units.

## Notes

### Competing Interest Statement

The authors have declared no competing interest.

### Funding

- NIH Common Fund: AI175078, AI181815, AI36920

